# Automatic Wake-Sleep Stages Classification using Electroencephalogram Instantaneous Frequency and Envelope Tracking

**DOI:** 10.1101/2020.05.13.092841

**Authors:** Mahdi Rahbar Alam, Reza Sameni

**Affiliations:** School of Electrical and Computer Engineering, Shiraz University, Shiraz, Iran.

**Keywords:** Automatic sleep staging, electroencephalogram, instantaneous frequency estimation, instantaneous envelope, LDA classification

## Abstract

**Background:** The study of cerebral activity during sleep using the electroencephalograph (EEG) is a major research field in neuroscience. Despite the rich literature in this field, the automatic and accurate categorization of wake-sleep stages remains an open problem.

**New Method:** A robust model-based Kalman filtering scheme is proposed for tracking the poles of a second order time-varying autoregressive model fitted over the EEG acquired during different wake/sleep stages. The pole angle/phase is regarded as the dominant frequency of the EEG spectrum (known as the instantaneous frequency in literature). The frequency resolution is improved by splitting the wide frequency band to subbands corresponding to well-known brain rhythms. Using recent findings in field of EEG phase/frequency tracking, the instantaneous envelope of the narrow-band signal’s analytic form is also tracked as a complementary feature.

**Results:** The minimal set of instantaneous frequency and envelope features is employed in three classification schemes, using training labels from R&k and AASM sleep scoring standards. The LDA classifier resulted in the highest performance using the proposed feature set.

**Comparison with Existing Methods:** The proposed method resulted in a higher mean decoding accuracy and a lower standard deviation on the entire dataset, as compared with state-of-the-art techniques.

**Conclusions:** The accurate tracking of the instantaneous frequency and envelope are highly informative for sleep stage scoring. The proposed method is shown to have additional applications, including the prediction of wake-sleep transition, which can be used for drowsiness detection from the EEG.

## 1. Introduction

Sleep is a state of unconsciousness, from which a person can be aroused. The neuronal activity of the brain dynamically varies throughout the states of wakefulness and different sleep stages. The electroencephalogram (EEG)— the most common means of studying the electrical activity of the brain— combined with other biological signals, such as the EOG, EMG, ECG, Spo2, etc. are used for polysomnography (PSG), which is currently the major tool for sleep studying.

A large number of human disorders affect the sleep quality. After pain, sleep disturbances are considered as the second most frequent indicator of human illness [1]. To date, there are more than seventy known sleep disorders. The most common sleep disorders include insomnia, narcolepsy, sleep apnea and restless legs syndrome (RLS), most of which can be managed effectively once they are correctly diagnosed [2].

Sleep stages recognition is a prerequisite for sleep disorders diagnosis; since certain disorders are known to occur in particular sleep stages. Moreover, the length and frequency of different sleep stages and cycles are prominent features in sleep study.

Sleep stages are generally divided into *rapid eye movement* (REM) and *non-REM* (NREM) states, each characterized with distinct physiological, neurological and psychological features. NREM is followed by REM sleep in which the brain activity increases up to wakefulness state. For this reason, REM is also called paradoxical or desynchronized sleep, due to physiological similarities (including brain electrical waves properties) to waking state. NREM is further subdivided into four stages of S1 (very light sleep), S2 (light sleep), S3 (deep sleep), and S4 (very deep sleep) based on the 1968 Rechtschaffen and Kales (R&K) sleep scoring standard [3]. The R&K was the gold standard until 2009, with the introduction of new sleep scoring guidelines by the American Academy of Sleep Medicine (known as the AASM standard), which made some major modification in scoring rules besides some data acquisition improvements [4]. In the AASM standard, the NREM stages S3 and S4 are combined into a single stage (called N3) known as slow wave sleep (SWS).

In practice, manual sleep scoring is done by visual inspection of PSG signals based on the noted scoring standards and guidelines. To date, many methods have been developed to automate the classification of wake-sleep stages from PSG or EEG alone. The numerous features used in sleep EEG analysis literature, include energy features, Fourier transform coefficients [5, 6], wavelet coefficients [7, 8, 9, 10], entropic features [6, 11], and fractal features [12]. The classification methods used in this area include discriminant analysis [12, 11], hidden Markov models [5, 13, 14], neural networks [8, 15, 10, 9, 16, 17, 18, 19], random forest [20, 11] and support vector machines [21, 22, 23, 24, 25].

Perhaps, among all the features the most common used for sleep EEG are the spectral features [26]. The EEG power spectral density (PSD) has been shown to be highly correlated with the different frequency bands of neurons activity (brain rhythms). Numerous methods have been used for EEG PSD estimation, which are generally based on the assumption of EEG stationarity during the estimation time interval. On the other hand, the EEG time interval length plays an important role in the time-frequency resolution, since the time and frequency resolution may not be simultaneously improved beyond the Heisenberg-Gabor lower bound [27]. Although the PSD provides an estimate of the per-Hertz energy distribution of the EEG over a given time interval, for sleep stages studies, we are more interested in the dynamics of the frequency components, which requires the tracking of the dominant instantaneous frequency of the EEG over time, corresponding to the neuronal oscillatory behavior, which may not be found in PSD analysis.

The application of *autoregressive moving-average* (ARMA) model root tracking using an RLS algorithm for EEG analysis was first introduced in [28]. It has been proposed that the pole displacement of an *autoregressive* (AR) model fitted over a nonstationary EEG can have physiological interpretations [29, 28, 30]. In [31] it was shown that the IF is the most correlated feature with variations in level of consciousness in the problem of depth of anesthesia (DOA) detection.

In this work, the *instantaneous frequency* and *instantaneous envelope* of EEG acquired during wake and sleep stages are estimated from various frequency sub-bands as spectral and energy features for automatic sleep stage scoring. The proposed method for IF and IE tracking is based on tracking the root loci of an AR model fitted over the EEG, as previously proposed in [32]. The major objective of this study is to show that by using conventional classifiers, a minimal (yet highly informative) feature set and an accurate Kalman filtering instantaneous envelope/frequency tracking scheme, it is possible to develop an accurate and low-complexity sleep scoring system.

## 2. Robust Instantaneous Envelope & Frequency Estimation

In this section, the methodology used to extract instantaneous envelope and frequency features from individual channels and frequency bands of raw EEG is explained in details.

### 2.1. The Concept of Instantaneous Envelope & Frequency

In non-stationary signals such as the EEG, the amplitude and frequency both vary in time, and it is useful to have an estimate of their instantaneous variations to characterize their non-stationarity. The instantaneous envelope and instantaneous frequency of a signal can be uniquely extracted from its *analytic form* [33]. Let *y_a_*(*n*) be the analytic form of a narrow-band discrete-time signal *y*(*n*):

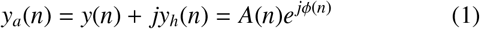

where *y_h_*(*n*) represents the Hilbert transform of *y*(*n*), *A*(*n*) is the analytical form modulus, known as the *instantaneous envelope* (IE), and *ϕ*(*n*) is the *instantaneous phase* (IP) of *y_a_*(*n*).

The *instantaneous frequency* (IF) is a scaled time derivative of the IP. For discrete-time signals and in low frequencies (relative to the sampling frequency), the IF can be approximated by the normalized first order difference of the IP mapped onto [0, 2*π*):

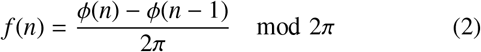

The estimation of EEG IF and IE is highly susceptible to the level of background cerebral activity. It was recently shown how the estimation of these parameters can be performed within a robust statistical framework [34]. Herein, we utilize a robust extension of the IF estimation algorithm utilized in [32, 30], by employing a variable dynamic model and considering the impact of tracking the combination of IF and IE features for the application of interest.

### 2.2. Bandpass Filtering and Decimation

The concept of instantaneous frequency is (implicitly) based on the assumption of a dominant frequency peak at each time instant [35]. However, for time-varying multi-component frequency signals such as the EEG, IF extraction is a challenge. to overcome this limitation, band-pass filtering prior to frequency domain feature extraction is a common approach. Considering a particular brain rhythm in the frequency band (*f*_L_, *f*_H_), the central frequency (*f*_C_) is equal to 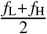. The maximum frequency (the Nyquist rate) is *f*_N_ = 2 *f*_H_, which is the minimum required sampling frequency in which the frequency information is preserved while sampling. At this rate, some morphological information of the EEG maybe lost, which is not important for automatic EEG signal processing where the aim is to extract a feature vector, rather than visual inspection by a clinician. Therefore, down-sampling the raw signal to this rate not only eliminates the unwanted frequencies for band-pass filtering; but also significantly reduces the processing complexity for long EEG recordings.

Bandpass filter is performed by applying a zero-phase forward-backward FIR filter, with lower and upper cutoff frequencies *f*_L_ and *f*_H_, respectively. IF estimation is applied at the output of the bandpass filter.

Frequency tracking is performed simultaneously in different EEG frequency bands (delta, theta, alpha and beta), to monitor the IF variations in the major EEG subbands simultaneously.

### 2.3. Amplitude Normalization

Amplitude normalization has been applied to the EEG modulus in [32], to eliminate the frequency estimation bias. It has been reported that the procedure has significant impact, practically in accurate estimation of modulus and phase of TVAR filter response. Herein, the following procedure is used for amplitude normalization, to make the estimation of the Averaged Variances Ratio (AVR) and the KF tuning parameters independent from the signal modulus:

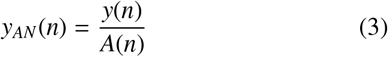

### 2.4. Time-Varying Autoregressive Model for IF Estimation

To date, various techniques have been applied to non-stationary signals for IF estimation. They can be generally categorized in two classes of parametric and non-parametric methods. Short Time Fourier Transform (STFT) [36, 37], Hilbert Transform [38] and wavelet based algorithms [27] are among the most common non-parametric methods, where the IF estimation is a tradeoff between the time and frequency resolution (according to the Heisenberg-Gabor time-frequency uncertainty principle [27]). On the other hand, in parametric methods– depending on the signal complexity– a linear or nonlinear model is considered for the signal. Autoregressive and Autoregressive Moving Average (ARMA) are two linear parametric models, which have been used for biomedical signal modeling. Time Varying Autoregressive (TVAR) model is a variable-parameter version of the AR model, which is appropriate for non-stationary signals such as the EEG, with time variant spectral characteristics [39].

In the AR model, a stationary discrete-time signal *y*(*n*) is expressed as the output of a linear time-invariant system with white noise input:

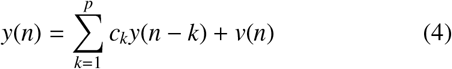

where *c_k_* ∈ {*c*_1_, ···, *c_p_*} are constant model coefficients, *v*(*n*) ~ (0, *R_n_*) is zero-mean white Gaussian noise (WGN), and *p* is the order of the AR model.

Accordingly, the AR process can be obtained by applying WGN to the following filter:

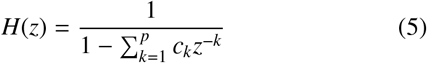

where, the pole configuration (including the number, radius and angle of poles) of *H*(*z*) determine the spectral behavior of the generated stochastic process. Accordingly, the number of oscillatory components depends on *p*, the order of the AR model. For example, by selecting a second-order AR model, denoted by AR(2), one assumes a single dominant frequency component. In general, depending on the pole loci, AR(2) can be a low-pass, band-pass or high-pass filter. The filter bandwidth *bw* and its center frequency *ω*_0_ are determined by the pole radius *r* and the pole angle *ϕ*, respectively.

The relation between the location of AR(2) complex conjugate poles in the z-plane and the parameters of the corresponding filter (*bw*,*ω*_0_) is illustrated in Fig. 1. The magnitude response *H*(*e^jω^*) shows that as the pole radius approaches the unit circle, the filter bandwidth (*bw*) decreases until reaching a narrow-band filter which exhibits a resonance at the center radian frequency *ω*_0_.

**Figure 1:**
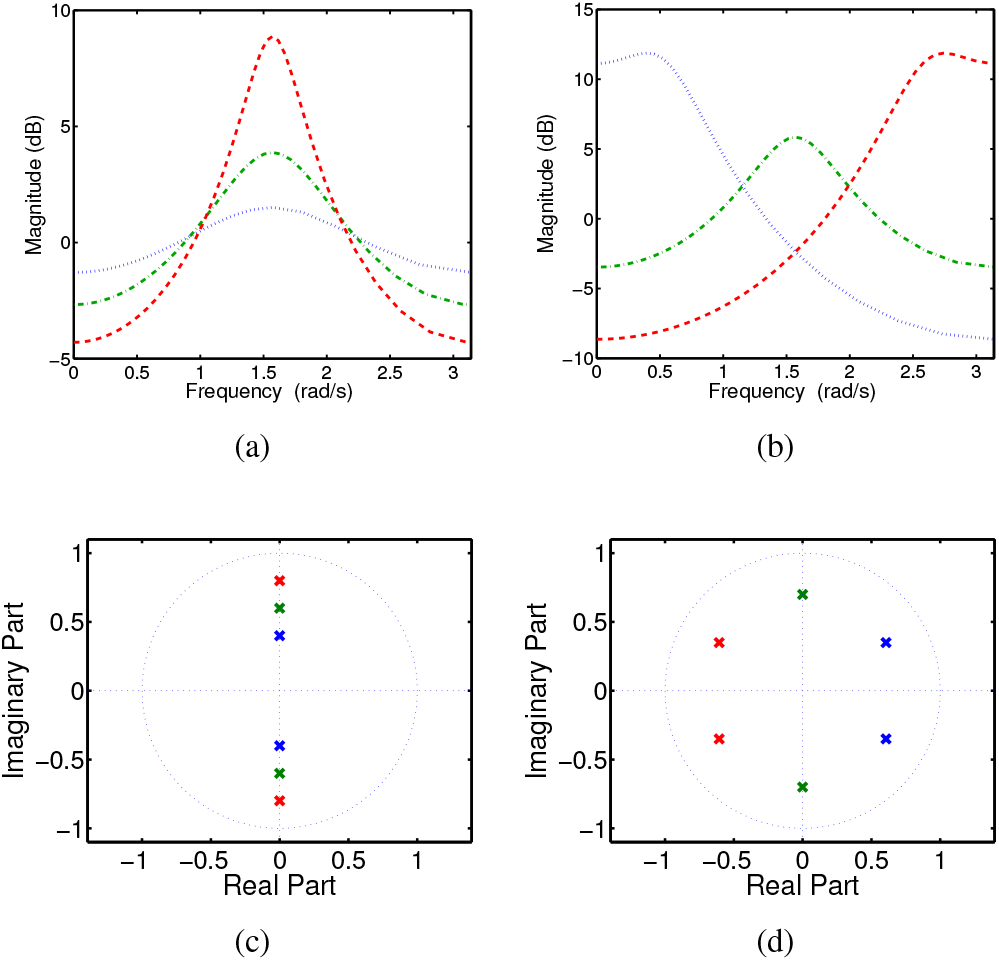
Autoregressive model poles location vs. AR(2) filter characteristics

In non-stationary signals such as the EEG, the frequency response *H*(*z*) is time-variant. Nevertheless, the displacement of the poles (and the variations of the corresponding transfer function) are rather slow from one sample to another. The frequency response of a time-varying AR (TVAR) of order two, denoted as TVAR(2), is as follows:

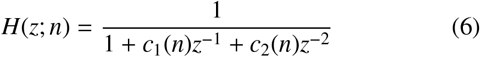

The magnitude of poles is related to the stability condition of the resulting stochastic process [40]. It can be shown that the following condition guarantees the stability of the filter output process:

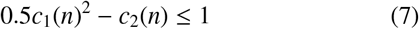

In the EEG, the time-varying center radian frequency *ω*_0_(*n*) has been attributed to the frequency at which the majority of cortical neurons in the recording lead vicinity oscillate in synchrony [30]. This is herein referred to as the *instantaneous frequency* (IF(*n*)) of the EEG, in Hz. The pole angle *ϕ*_0_(*n*) corresponds to the radial resonance frequency *ω*_0_(*n*) via the relation:

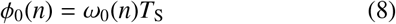

where *T*_S_ is the sampling period of the signal under analysis and *ϕ*_0_(*n*) ∈ [0, 2*π*]. IF(*n*) is next obtained as follows:

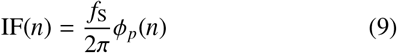

### 2.5. TVAR Model Parameter Tracking using Kalman Filter

The Kalman Filter (KF) is one of the adaptive approaches, which have been used in the literature for TVAR coefficients tracking and estimation [41, 42, 32]. The linear KF with Gaussian noise provides an optimal estimate, in the sense of minimum mean square error [43]. These conditions hold very well for EEG signals with a TVAR model.

Considering the coefficients vector **c**_*n*_ = (*c*_1_(*n*), *c*_2_(*n*))*T* as the system’s state vector, TVAR(2) can be represented in state-space by the two following equations, which are required for the KF design.

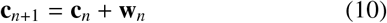

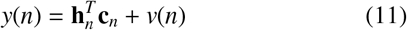

where *y*(*n*) is the observed EEG, **w**_*n*_ = (*w*_1_(*n*), *w*_2_(*n*))^*T*^ is considered as zero-mean white Gaussian *process noise* with covariance matrix

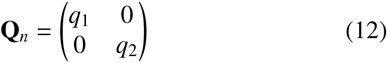

and *v*(*n*) is a scalar zero-mean white Gaussian *observation noise* with variance *r_n_*. Also, **h**_*n*_ = (*y*(*n* − 1), *y*(*n* − 2))^*T*^ is the observation transformation vector consisting of the two previous observations.

According to the state evolution equation (10), the coefficient vector evolution has been assumed to follow a first-order AR model, in the form of a Random Walk or Wiener process [44], which is a common assumption when no other priors are available about the state variations [45, 46].

The forward KF equations for sequential estimation of the state vector are as follows:

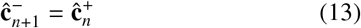

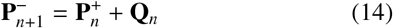

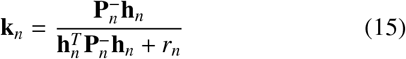

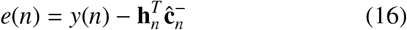

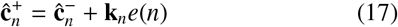

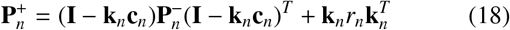

where **P**_*n*_ ∈ ℝ^2×2^ is the estimation covariance matrix, **I** is the two-by-two identity matrix, **k**_*n*_ is the *Kalman gain* vector and *e*(*n*) is the error in observation prediction, well-known as the *innovation signal*. In all equations the superscripts − and + refer to the estimation of the corresponding quantity before and after observation arrival, respectively.

Equation (18) is the stabilized form (known as the Joseph form [47]) for updating the state vector covariance matrix, which guarantees a positive semi-definite covariance update.

The recursion is initialized by 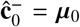 and 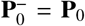 where µ_0_ = (*µ*_1_, *µ*_2_)^*T*^ and **P**_0_ ∈ ℝ^2×2^ are presumed mean and covariance of the initial Gaussian state vector 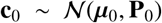. The initial state vector can be set by applying the Yule-Walker method on the entire signal offline.

The KF is causal, as it uses the two previous observations to estimate the current state. Better performance may be obtained if one uses a Kalman Smoother (KS). The KS basically consists of a forward KF followed by a backward recursive smoothing stage. Depending on the smoothing strategy, smoothing algorithms are usually classified into fixed-lag or fixed-interval smoothers [48, 49]. Herein, we use a fixed-interval KS for offline sleep staging. In this scheme, having the forward estimated states and their covariance matrix from sample *n* = 1 to *n* = *N*, the backward estimation process is applied recursively for *n* = *N* − 1 to *n* = 1 as follows:

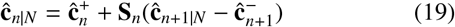

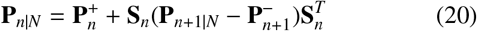

where

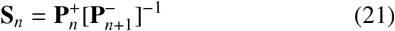

In these equations *n|N* denotes *n*-th smoothed version of the state vector or covariance matrix, using *N* samples of the same quantity. This implementation of the KS is known as the Rauch-Tung-Striebel two-pass KS algorithm^1^.

### 2.6. KF Parameter Tuning

It can be shown that for the dynamic model (10) and (11), the KF equations are only a function of the ration *γ* = *Q/R* [50, 48]. Therefore, one only needs to tune *γ* instead of both parameters *Q* and *R*. In order to set this ration, we hereby employ a monitoring criterion coined as the *averaged variances ratio* (AVR), which has been previously used in [49, 51] for monitoring the fidelity of the KF and updating the values of the KF covariances. AVR is defined as the variance of the KF innovation signal divided by the KF estimated variance over a sliding window of length *L*. Accordingly, for the *i*th estimation, the AVR criterion is:

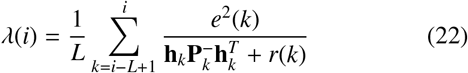

If the KF parameters (*γ* in our case) are selected correctly, one should have *λ*(*i*) ≈ 1. In practice the fluctuation of *λ*(*i*) between 0.5 and 2 with an average of 1 is acceptable [50]. In the current case, *γ* has been selected such that *λ*(*i*) fluctuates around 1 in the awake state. In real sleep studies, this procedure can be done per subject by a short learning process before sleep.

## 3. Simulation on Synthetic Signal

In order to show the problem of robust IF estimation, the following AM-FM sinusoidal signal is used for illustration:

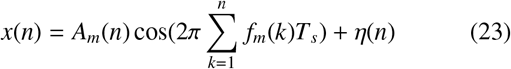

where *A_m_*(*n*) = 1.2 sin(2*π* × 0.75*n* + *π/*4) + 0.6 sin(2*π* × 2.5*n*) is the instantaneous envelope, *f_m_*(*n*) = 2.5 sin(2*π* × *n*) + 10.5 is the time-varying frequency (corresponding to the alpha band) and *η*(*n*) is a zero-mean white Gaussian noise with variance adjusted to set the signal-to-noise ratio (SNR) of *x*(*n*) to 20 dB. The signal is simulated at a sampling frequency *f_s_* =300 Hz.

Theoretical and practical challenges of IP and IF estimation were detailed in [34]. Abrupt random phase and frequency jumps were shown to occur more frequently in low analytical signal envelopes. This effect (the randomization of the estimated IF using conventional Hilbert transform methods in presence and lack of a BPF corresponding to alpha wave bandwidth have been illustrated in Figs. 2(c) and 2(b), respectively. Although band-pass filtering reduce the effect of out of range frequencies, but is not enough for a reliable IF extraction. The performance of IF estimation using two methods of Hilbert transform with band-pass filtering and the robust method is also compared in figure 2(c). By proper choice of Kalman filter/smoother parameters, the performance of this method is much more accurate in low analytical signal envelopes, despite some inevitable deflections around the true IF. This example motivates the temporal tracking of the IE and IF for a more robust estimation of EEG temporal frequency variations.

**Figure 2:**
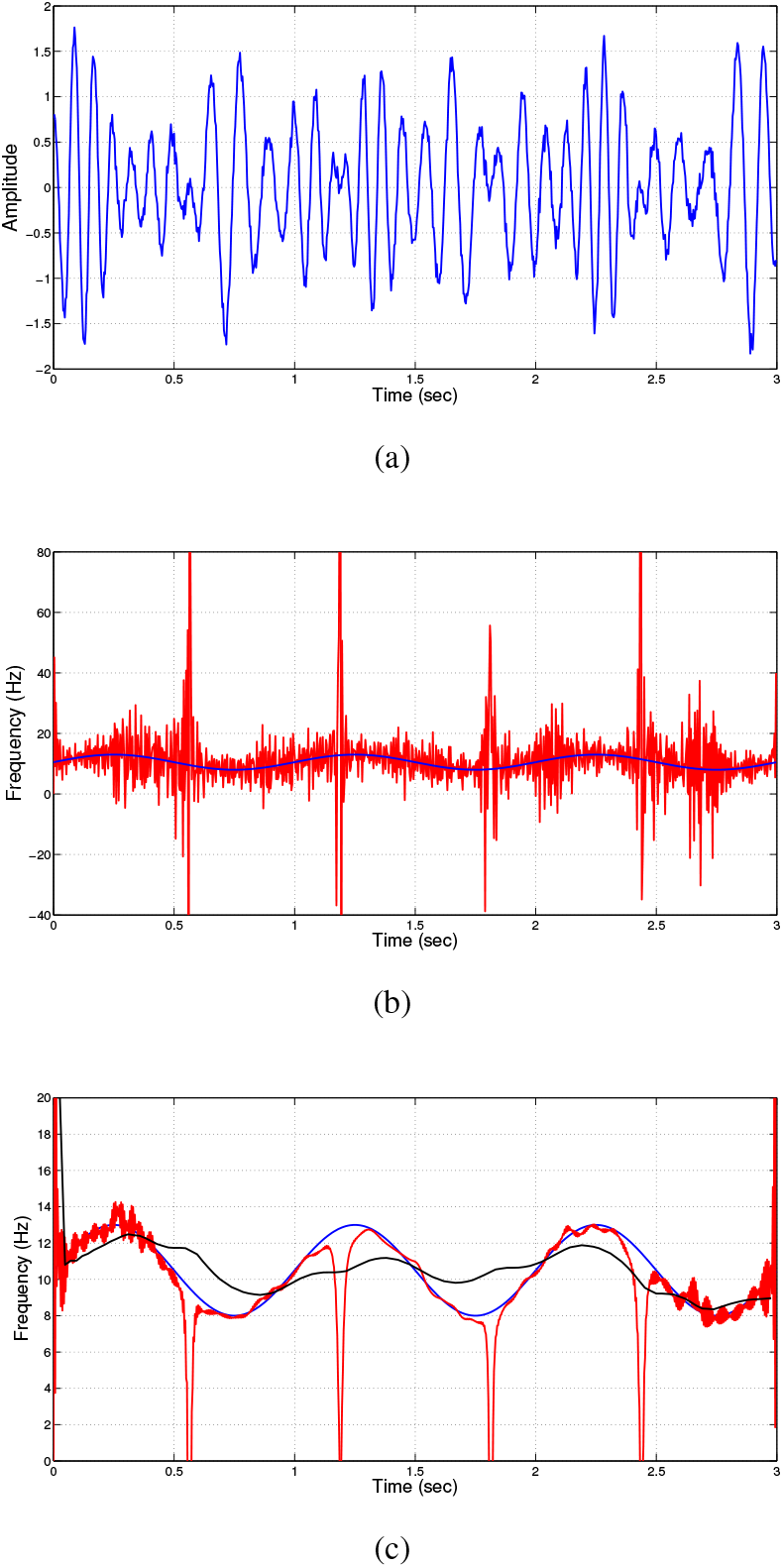
Simulated data: (a) a synthetic AM-FM noisy signal with SNR=20 dB; (b) The time-varying true IF (blue line) and the IF obtained via Hilbert Transform (red line); (c) The true IF (blue line); the instantaneous frequency obtained via band-passed Hilbert Transform (red line); the IF estimated via the KS (proposed algorithm) (black line)

## 4. Case Study

### 4.1. Dataset

In this study, two EEG channels, Fpz–Cz and Pz–Oz, are taken from the Sleep-EDF database, containing PSG recording described in [52]. According to this, The selected dataset contained of 39 sleep cassette PSG recorded from 20 healthy subjects including ten males and ten females within an age range of 25 to 34 years old, together with their *hypnogram* (expert annotations of sleep stages). The PSG of each subject was sampled at 100 Hz in two successive day-night periods (except for fourteenth subjects with only one night data acquisition) and the hypnogram index was reported for 30 s time intervals. The hypnogram labels consist of W (wakefulness state), 1, 2, 3, 4 (stages S1 to S4 of non-REM sleep), R (REM sleep) and M (movement time). Since the movement time and un-scored data segments were negligible as compared to the labeled parts, they were not taken into account and were omitted from analysis. Note that “movement times” have been eliminated in the more recent AASM standard [4]. All hypnograms were manually scored by well-trained technicians according to the Rechtschaffen and Kales manual [3]; but based on Fpz–Cz/Pz–Oz EEGs instead of C4-A1/C3-A2 EEGs, as suggested by [53].

The PSGs were recorded from the subjects during normal activities in both awake and sleep states. Since the data has been acquired mostly in the awake state, with small proportions of stages S1 and S4, we are dealing with an unbalanced dataset, which is an important factor for the classifier.

In order to examine the adaptability of the classification algorithms with the AASM standard, stages 3 and 4 were merged into slow wave sleep (SWS). Nevertheless, the other AASM parameters could not be fulfilled using the current dataset, due to the differences in scoring rules as comparing with the R&K standard.

### 4.2. Spectrum Analysis

It has been previously reported that the changes in consciousness level is proportional to the energy level variations of different frequency bands [31]. For illustration, this point is shown in the spectrogram of a sample ten-hour wakefulness and sleep EEG, acquired from the Fpz–Cz channel, together with its corresponding hypnogram, in Fig. 3.

**Figure 3:**
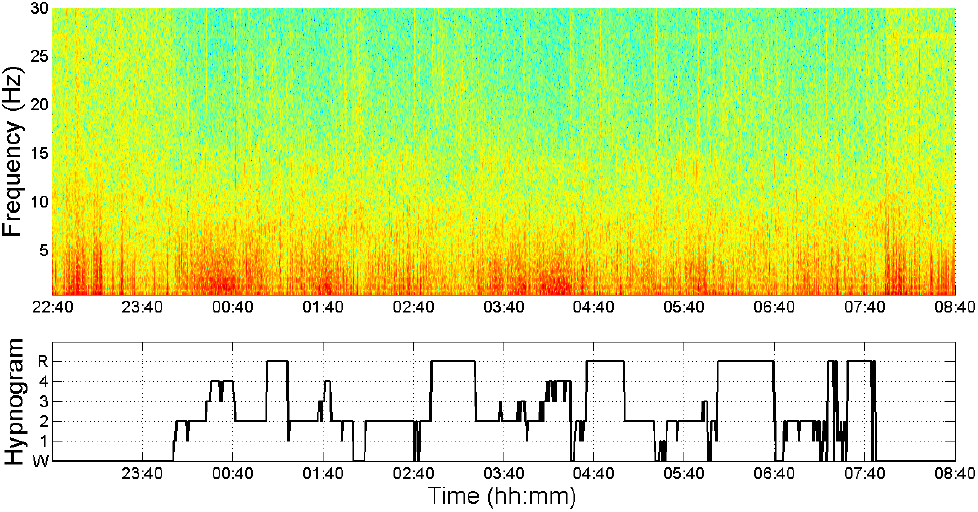
Spectrogram (top) and hypnogram (bottom) of different brain rhythms in wakefulness and sleep states for a ten-hour sample EEG

The wide-band spectrogram in Fig. 3 reveals that the sleep state is correlated with the energy variations of alpha and beta rhythms. In order to validate this observation more rigorously, we divide the wide-band frequency range of the EEG, into the sub-bands corresponding to common brain rhythms locating between 0.5–30.0 Hz. Therefore, the aforementioned IP/IF extraction algorithm is applied simultaneously on four parallel bands: *δ* (0.5–4.0 Hz), *θ* (4.0–7.5 Hz), *α* (8.0–13.0 Hz), and *β* (14.0–30.0 Hz).

### 4.3. Feature Extraction

The proposed algorithm has been applied on the entire dataset over all the four frequency bands detailed in the previous subsection, in order to extract the features of IE and IF, without any prior judgment about their efficiency in different bandwidths. The following procedure was implemented on both Fpz–Cz and Pz–Oz EEG channels separately. Figure 4 gives an overview of the feature extraction algorithm.

**Figure 4:**
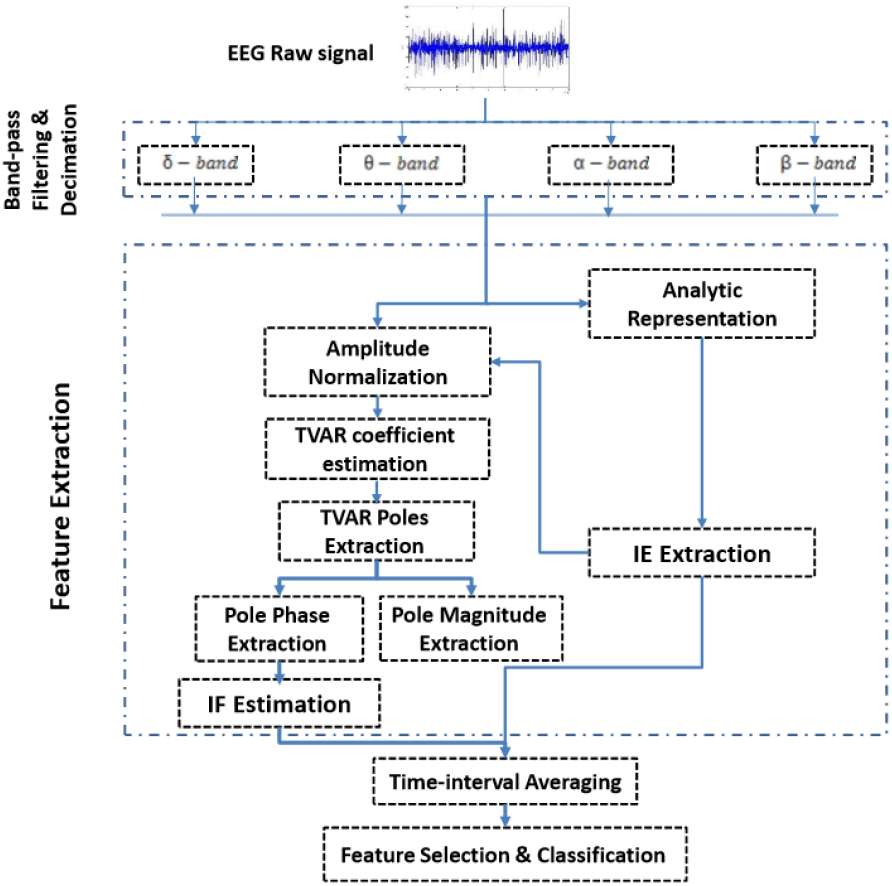
Feature extraction block diagram

According to figure 4:

1. In a pre-processing stage, the raw signal is decimated and passed through four parallel band-pass filters with bandwidth ranges corresponding to the *δ*, *θ*, *α* and *β* brain rhythms.
2. The bandpass signals are represented in analytic form (as in 1) and their modulus are extracted as the *instantaneous envelope* (IE).
3. The bandpass signals are normalized by dividing them by their IE.
4. The *instantaneous frequency* (IF) of the normalized bandpass signals are estimated using the robust algorithm considering the detailed implementation issues described in Section 2.
5. Stages 2 and 4 (above) result in eight IF & IE feature vectors (time series) per channel, with different lengths and sampling frequencies. Since, the hypnogram labels are available for 30 s intervals, *time-interval averaging* is used to average the feature samples over 30 s time intervals.

The above procedure is implemented on both EEG channels for all subjects. The whole feature vectors together with their hypnogram labels are saved for post-processing feature selection and classification algorithms.

### 4.4. Feature Selection & Classification

The effectiveness of the proposed feature vector is evaluated by running a classification algorithm, using the hypnogram labels for the training and test phases.

The proposed feature extraction algorithm results in a sixteen element feature vector. Herein, we take a multistage feature selection (rather than a blind and naïe feature reduction), to assess the effectiveness of single features and their combinations. This procedure enables a better interpretation of the proposed feature vector and eventually permits the reduction of the feature vector length.

The choice of classifier is subjective. In order to compare the performance of different classifiers on the current problem, we have implemented three techniques: Linear Discriminant analysis (LDA), One-Against-All Support Vector Machine (OAA-SVM) and the Dendrogram-based Support Vector Machine (D-SVM). All three classifiers have been applied in a recent work on sleep stages classification [25]. For benchmarking, the D-SVM Matlab codes have been adopted from [25], which are available online. The Leave-One-Subject-Out (LOSO) procedure is used for cross-validation of all three classifiers. Accordingly, the training and test feature sets are selected based on the LOSO scheme, such that the classifier is trained over nineteen subjects and tested on the remaining one. The classification performance is evaluated by the well-known *confusion matrix* metric. In addition, in order to report a reliable overall accuracy per subject, the *decoding accuracy* (DA) is also used [25], which is the mean of diagonal elements of the confusion matrix.

Discriminant analysis is a common technique for data classification and dimensionality reduction. Simply put, it projects the data to a space with highest between-class and lowest within-class variances, to achieve maximal separability. A Gaussian distribution is assumed for the samples per class; hence, the class means and covariance matrices are the only required parameters. In linear discriminant analysis, which is used in this study, we assume that the covariance matrices of both classes are identical and only the class means (class centers) vary.

Support Vector Machine classifiers are inherently designed for binary classification problems. Nevertheless, multi-class extensions of SVMs, such as the One-Against-One (OAO) and the One-Against-All (OAA) are also very popular in practice. In the hereby implemented OAA-SVM, a Radial Basis Function (RBF) kernel it fitted over the training data per class. In the test phase, the data is evaluated for each class model with a prediction probability. At last, the test data is associated to the class with highest probability.

Employing the D-SVM (a decision tree multi-SVM classifier) for wake/sleep stages classification was proposed in [25]. In the D-SVM framework, a tree of ascending or descending hierarchical clusters (AHC or DHC) is produced, based on a measure of distance between the representatives of clusters (classes).

Herein, following [25], an ascending hierarchy (AHC) is employed. The mean value of classes is assigned to be the representative of clusters and the distance measure is chosen to be the Euclidean distance divided by the standard deviation of the cluster. This measure provides maximal discrimination between classes based on the Fisher criterion. The generated dendrogram is then used as the backbone for multi-class classification, with a distinct binary SVM assigned to each node [25].

## 5. Results

In this section the implementation results of the explained algorithm applied on the entire thirty-nine ensembles of the introduced dataset is presented as follows. First, the estimated features are visualized and then the certainty about the estimated IF is presented with the confidence interval measure. Next, a comparison between the two implemented IF estimation approaches from the distribution point of view is performed. Finally, the overall and subject-wise performance of the proposed algorithm is analyzed.

### 5.1. Visual Analysis

A typical sample of the estimated IF and IE in four brain rhythms together with the corresponding hypnogram is illustrated in Fig. 5. Due to the bandpass filtering procedure, all the estimated IFs are located in the corresponding bandwidth. By visual inspection, the frequency variations of the hypnogram are somehow correlated with the IF variations. Nevertheless, visual inspection of a single trial is insufficient for drawing a general conclusion. Herein, following the findings of [34], regarding the unreliability of the IF in low analytical signal envelopes, the IE is used as a complement for the IF.

**Figure 5:**
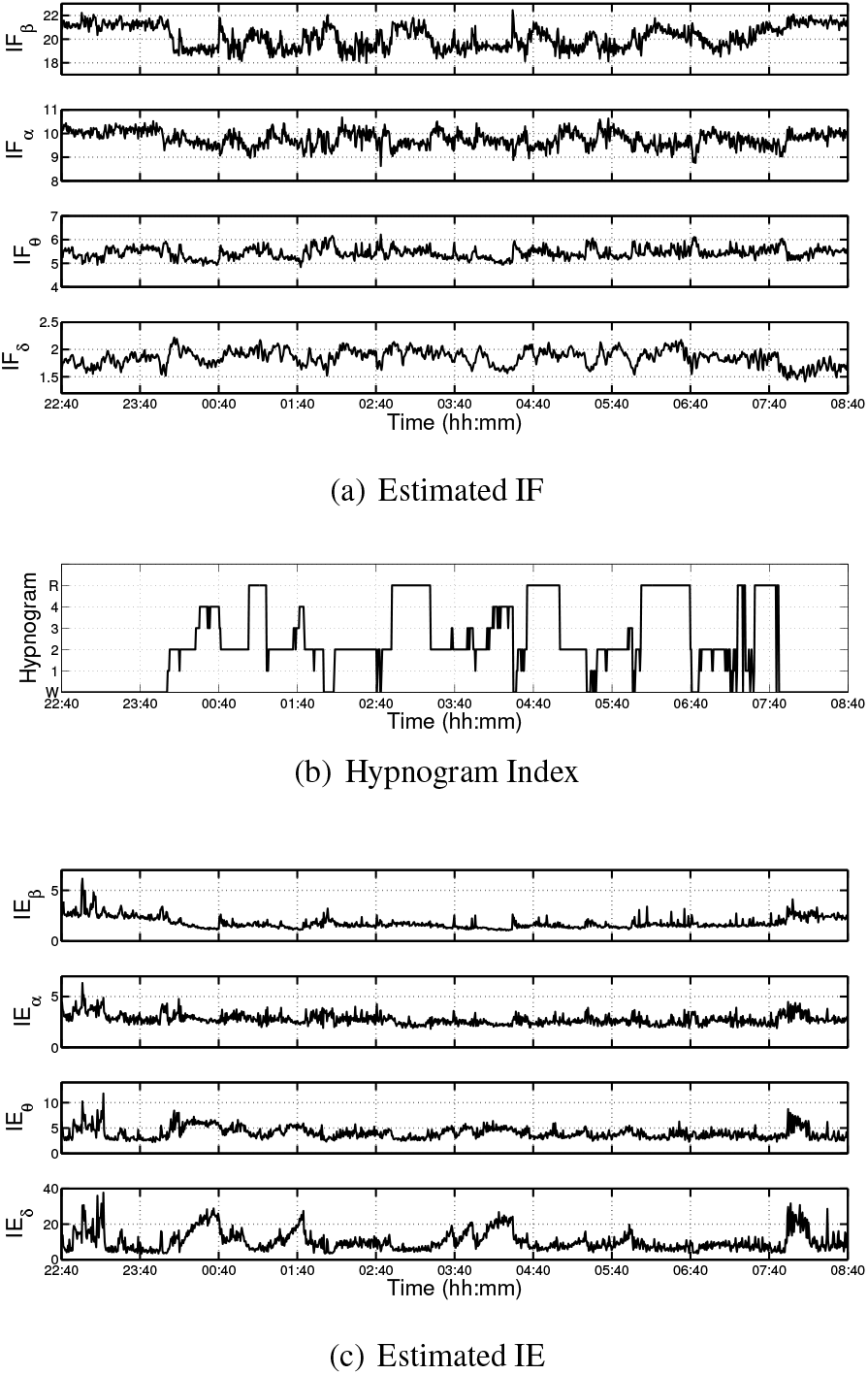
Visualization of the estimated IF and IE in different EEG bandwidths and the corresponding hypnogram index over the estimation interval

### 5.2. IF Estimate Confidence Interval

To obtain a confidence measure for the estimated IF values, we propose to monitor the lower and upper estimation bounds defined by the intervals [IF(*n*) − *σ*_IF_(*n*), IF(*n*) + *σ*_IF_(*n*)], where *σ*_IF_ is the standard deviation of the estimated values. For illustration, the estimated IF_*δ*_ tracked with its confidence interval and the magnitude of the corresponding AR model pole is illustrated in Fig. 6. Accordingly, as the pole magnitude approaches one (the unit circle), the uncertainty in the estimated IF decreases. It is known that for a narrow-band signal, when the generating model pole magnitudes approach the unit circle, the system output becomes more oscillatory. This means that in a narrow-banded EEG signal an *oscillatory instantaneous pole* could potentially have spontaneous or event-related physiological interpretations. In multichannel EEG analysis this measure may additionally result in the localization of oscillatory frequency generating sources.

**Figure 6:**
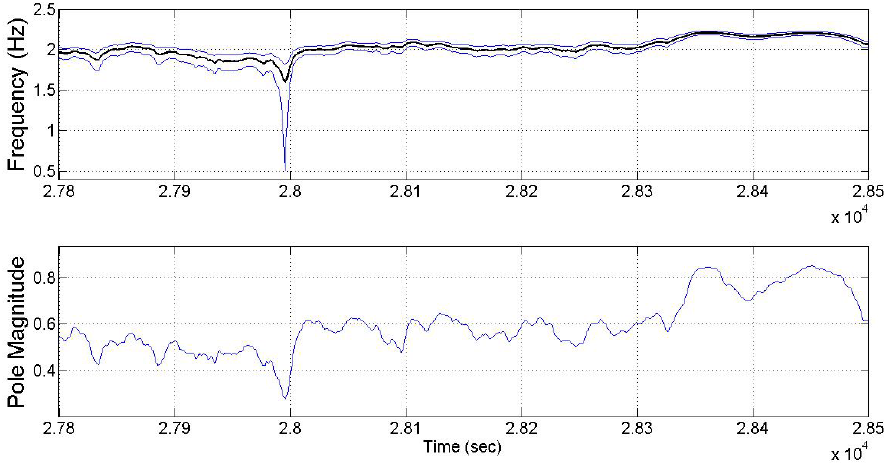
(Top) the estimated IF with its plus/minus standard deviation confidence intervals; (bottom) AR model pole magnitude

### 5.3. Comparison of Methods

In Section 2.5, the ability of KS in IF tracking was detailed. Herein, a comparison between the conventional Hilbert transform method and the KS approach for instantaneous frequency estimation is shown on real data.

Fig. 7 illustrates the sample distribution of IF_*β*_ for 2000 random samples per sleep stage belonging to one subject. The raw IF extracted by the Hilbert transform (the colorful points) and the Kalman Smoother methods (black dots) are compared, without any averaging. Apparently, the IF estimated by the Hilbert transform is not discriminative between the sleep stages. However, the IF distribution after applying the KS is even visually discriminative between the different classes (wake/sleep stages).

**Figure 7:**
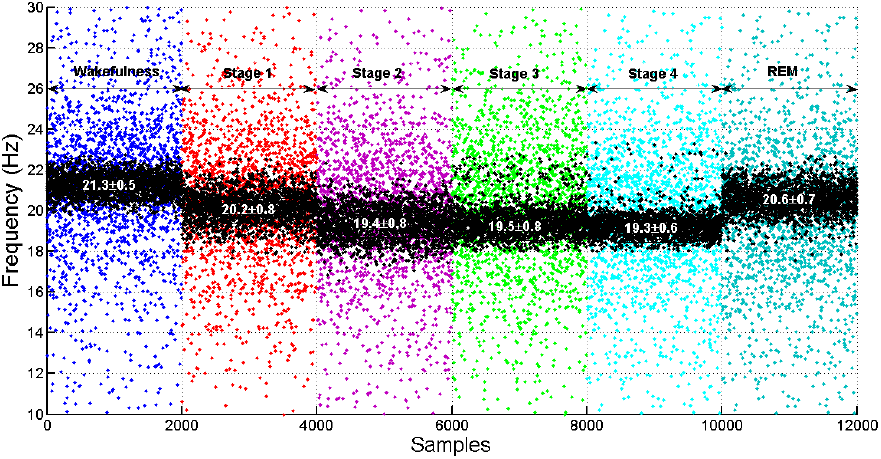
Sample distribution comparison of the instantaneous frequency of the beta wave for 2000 random samples per wake/sleep stages obtained via two methods. The colorful samples correspond to the Hilbert transform method and the black ones correspond to the Kalman smoother approach

In addition, according to Fig. 7, the mean value of the beta IF in the REM stage is close to the wakefulness frequency, which is in accordance to the physiological properties of the REM sleep.

### 5.4. Classification Results

According to the described feature selection strategy, an exhaustive search was carried out to find the best feature and combination of features using the LDA classifier. The results are shown in Table 1. Accordingly, if one seeks a single feature, IF_*β*_ has scored the best among all the proposed features for both channels (above 36% alone between five classes). If one selects more than a single feature, the best combination of features differs in the two channels. In the proposed scheme, at each step, among the remaining features, a single feature is added to the previous optimal feature subset of the previous step. As the feature vector size increases, the performance improves up to eight features listed in Table 1 (except when adding IE_*θ*_ of channel 2). Therefore, a subset of 15 features is selected for classification.

**Table 1:** Optimal Feature Combinations in Terms of Classification Accuracy

| # of Features | Channel 1 |  | Channel 2 |  | Channel 1&2 |
| --- | --- | --- | --- | --- | --- |
| | Best Combination | Mean DA±std | Best Combination | Mean DA±std | Mean DA±std |
| 1 | IF$_{\beta}$ | 36.53±1.97 | IF$_{\beta}$ | 38.58±2.11 | 44.90±6.08 |
| 2 | IF$_{\delta}$, IF$_{\beta}$ | 51.76±4.75 | IE$_{\delta}$, IF$_{\beta}$ | 47.43±6.67 | 60.64±6.36 |
| 3 | IE$_{\delta}$, IF$_{\delta}$, IF$_{\beta}$ | 60.63±5.35 | IE$_{\delta}$, IF$_{\alpha}$, IF$_{\beta}$ | 53.86±9.07 | 66.85±6.22 |
| 4 | IE$_{\delta}$, IE$_{\theta}$, IF$_{\delta}$, IF$_{\beta}$ | 62.95±5.81 | IE$_{\delta}$, IF$_{\delta}$, IF$_{\alpha}$, IF$_{\beta}$ | 57.50±8.13 | 69.33±5.82 |
| 5 | IE$_{\delta}$, IE$_{\theta}$, IE$_{\beta}$, IF$_{\delta}$, IF$_{\beta}$ | 65.11±5.76 | IE$_{\delta}$, IE$_{\beta}$, IF$_{\delta}$, IF$_{\alpha}$, IF$_{\beta}$ | 59.69±7.42 | 70.64±5.47 |
| 6 | IE$_{\delta}$, IE$_{\theta}$, IE$_{\beta}$, IF$_{\delta}$, IF$_{\theta}$, IF$_{\beta}$ | 66.88±6.18 | IE$_{\delta}$, IE$_{\beta}$, IF$_{\delta}$, IF$_{\theta}$, IF$_{\alpha}$, IF$_{\beta}$ | 61.24±7.23 | 73.02±5.19 |
| 7 | IE$_{\delta}$, IE$_{\theta}$, IE$_{\alpha}$, IE$_{\beta}$, IF$_{\delta}$, IF$_{\theta}$, IF$_{\beta}$ | 67.84±5.95 | IE$_{\delta}$, IE$_{\alpha}$, IE$_{\beta}$, IF$_{\delta}$, IF$_{\theta}$, IF$_{\alpha}$, IF$_{\beta}$ | 61.86±6.81 | 73.46±4.62 |
| 8 | IE$_{\delta}$, IE$_{\theta}$, IE$_{\alpha}$, IE$_{\beta}$, IF$_{\delta}$, IF$_{\theta}$, IF$_{\alpha}$, IF$_{\beta}$ | 68.42±6.10 | IE$_{\delta}$, IE$_{\theta}$, IE$_{\alpha}$, IE$_{\beta}$, IF$_{\delta}$, IF$_{\theta}$, IF$_{\alpha}$, IF$_{\beta}$ | 61.68±7.15 | 72.61±5.30 |

The classification training phase of DSVM forms the decision trees for the two sleep scoring standards, as shown in Fig. 8. The hierarchical clustering indicates high proximity of clusters (classes) corresponding to stages S1, REM and stages S3, S4 in the R&K standard. We have the same observation regarding stages {S1, REM} in the AASM standard.

**Figure 8:**
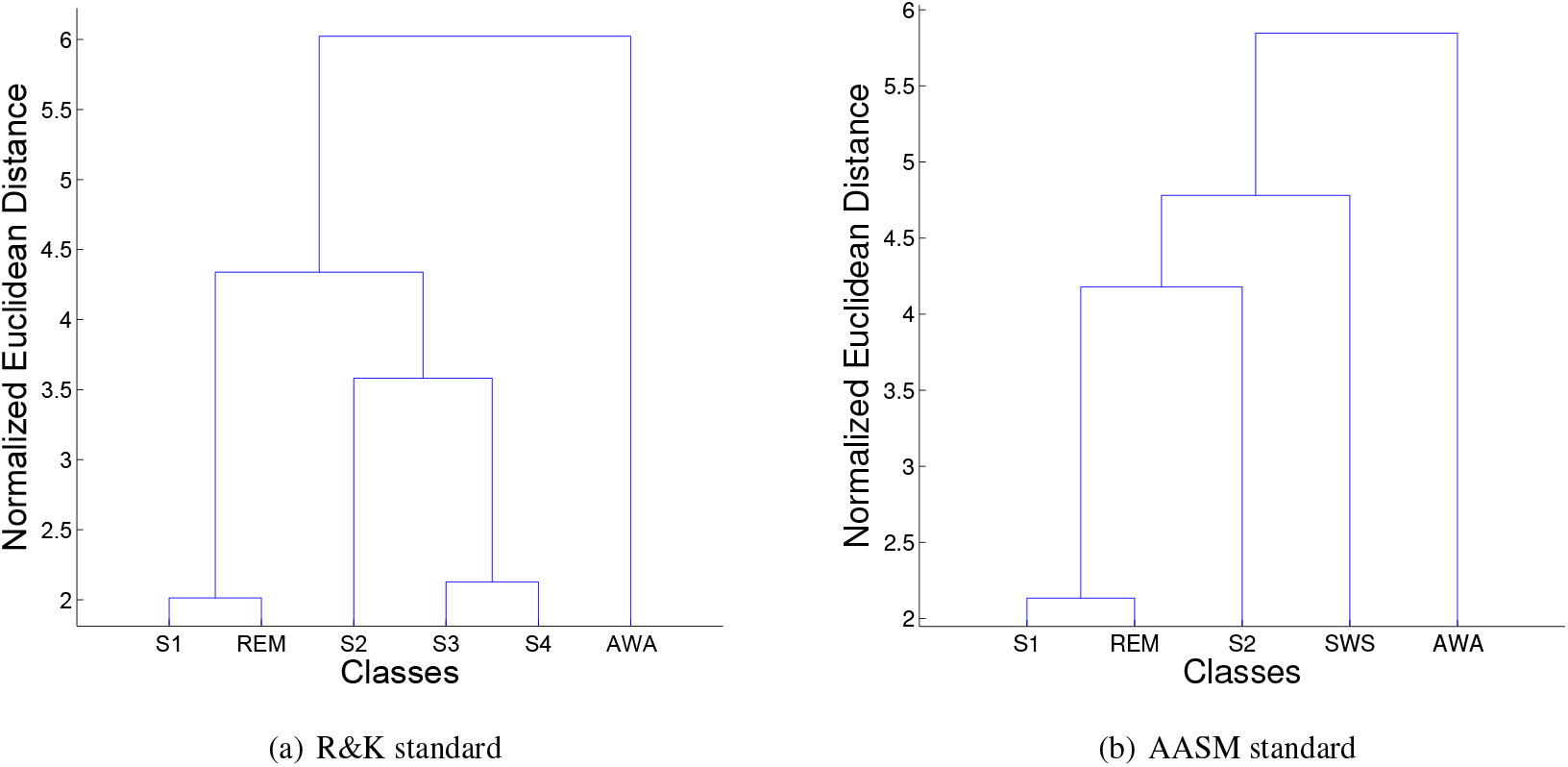
The decision trees formed from hierarchical clustering of DSVM training phase for two sleep scoring standards

The results of classification by the three described methods on the whole dataset are represented in confusion matrices of Fig. 9, for both sleep scoring standards. The results are in accordance with the pre-described hierarchical clustering structure in Fig. 8. Overall, the wakefulness state was classified with the highest accuracy rate. The first stage of sleep could be hardly distinguishable from the wake and/or REM stage using this method. Stage S2 was detected with acceptable accuracies, depending on the classification methodology. The notable point about stages S3 and S4 in the R&K standard is their interchangeable mis-classification. This problem was mostly eliminated by their combination in the AASM sleep scoring standard. Finally, the REM stage has the best accuracy in LDA method.

**Figure 9:**
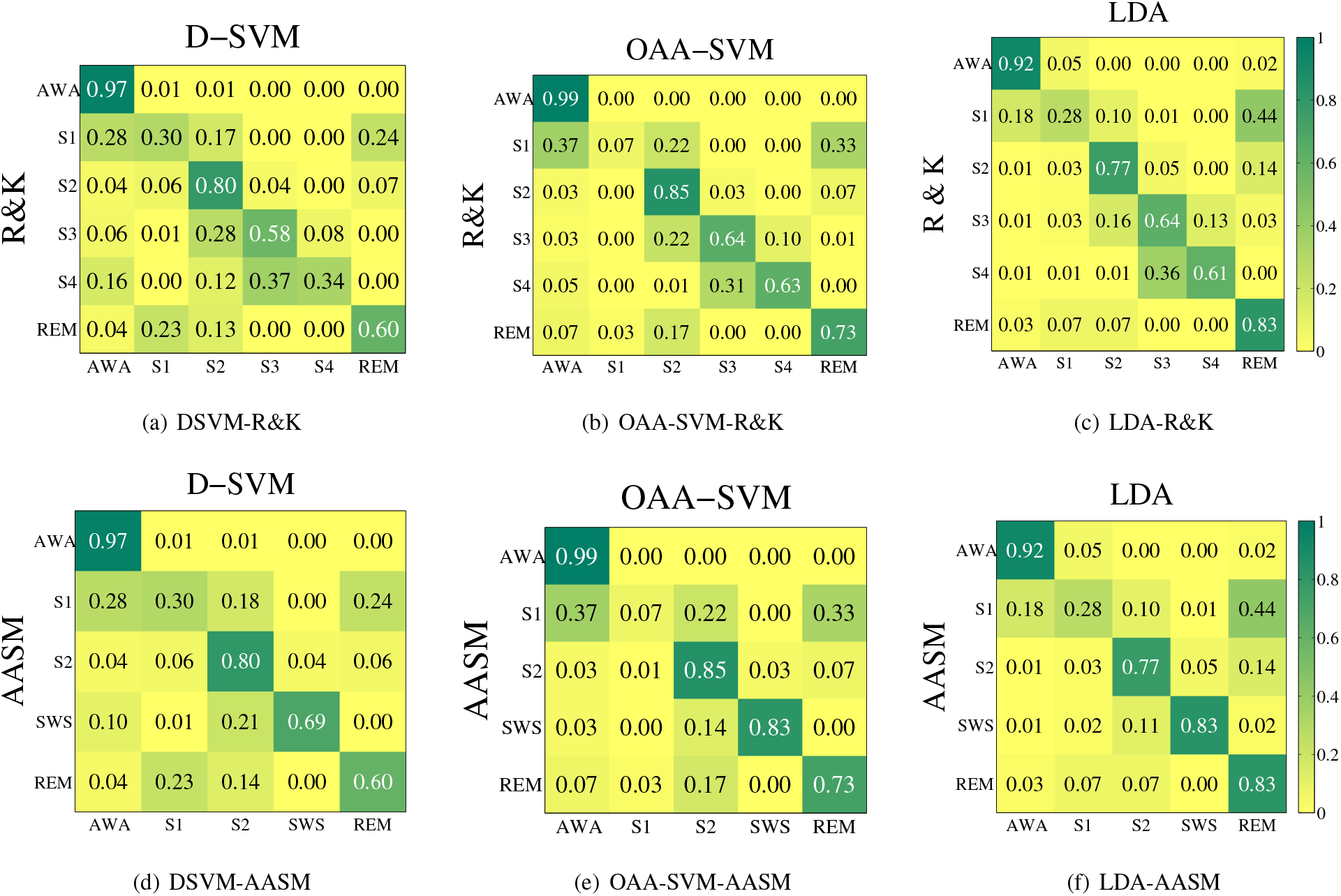
Confusion matrices for DSVM, OAA-SVM and LDA classification methods based on the R&K and AASM sleep scoring standards

According to the confusion matrices of the three different classification methods, it can be inferred that LDA has a higher overall mean accuracy and reliability as compared with other classifiers, using the proposed feature vector.

### 5.5. Individual performances

The performance of the LDA classifier can be better observed in the subject-wise results. Employing the Leave-One-Subject-Out strategy gives two test EEG ensembles for each subject, being trained by nineteen other unseen subjects (including 38 ensembles). The decoding accuracy (mean of individual class accuracies) was used— as in literature— for evaluation. The bar graph of Fig. 10 indicates this measure for all twenty subjects. The highest DA was 80% for subject 16 and the lowest was 63% for subject 5. The overall mean and standard deviation were 72.84% and 5.21%, respectively. The results have clearly outperformed the mean DA std=64.75 10.37% of the previous research [25].

**Figure 10:**
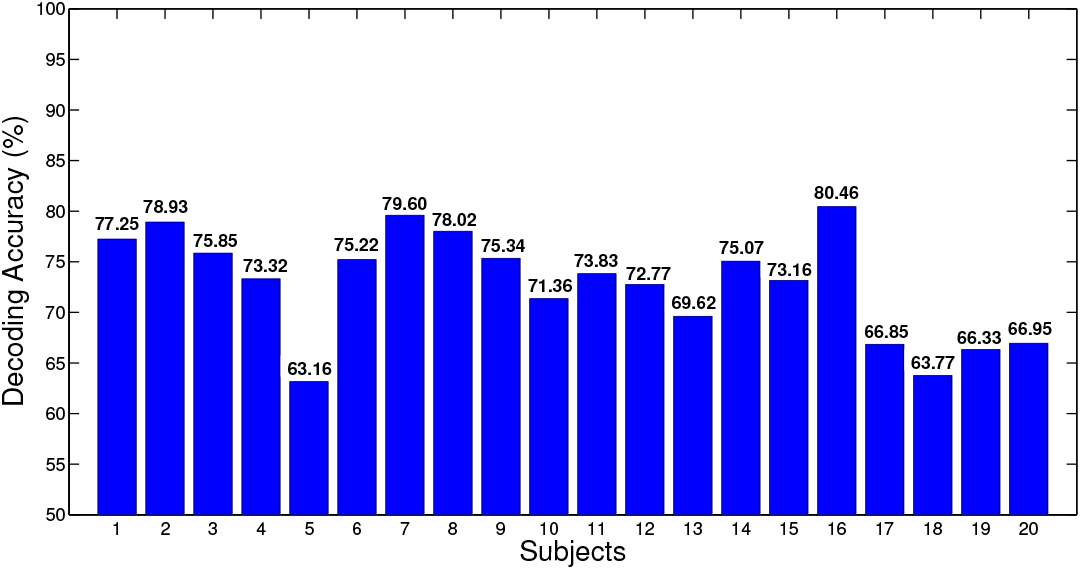
Decoding Accuracy (DA) for individual subjects using LDA classification

## 6. Discussion

The proposed methodology has various implications in the field of consciousness analysis and may be extended from different aspects in future studies. Two major points are detailed below.

### 6.1. Drowsiness Detection

Besides sleep stage analysis, a potential application for the proposed method is for drowsiness detection. Let us consider the AR model pole-magnitude variations along the hypnogram. By visual inspection of multiple subjects, the *α*-wave AR model pole-magnitude extracted from the occipital region channel has particular and rather exclusive behaviors during awake-sleep transition. For proof of concept, a typical case of the *α*-wave pole-magnitude extracted from the Pz–Oz channel together with the corresponding hypnogram has been plotted in Fig. 11. Accordingly, the foregoing pole amplitude approaches one in a short time prior to the onset of the first sleep stage and it returns to its normal value after the sleep stage begins. The time interval of this transient oscillatory behavior is of course subjective. The same behavior is observable during sleep-wake transition.

**Figure 11:**
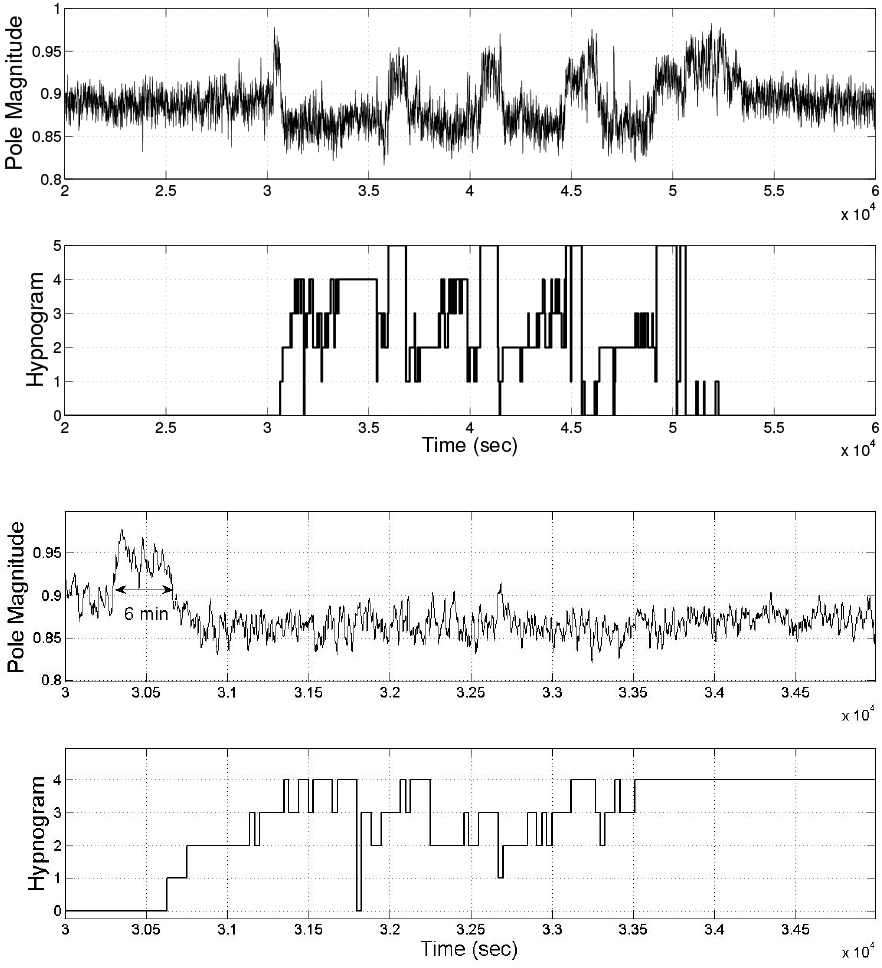
(Top two rows) Instantaneous pole magnitude of the second channel corresponding in the *α* band and the corresponding hypnogram index; (bottom two rows) The magnified plot of the top two rows highlighting the transition from wakefulness to sleep

Considering the dominant appearance of the *α* rhythm during sleep [54], the hereby observed phenomenon can be considered as a transition between wake and sleep stages, or *drowsiness*. The monitoring of drowsiness has significant impact on alertness detection for real-world applications, such as drivers, pilots, control center operators, etc. Unfortunately, standard datasets such as the ones used in this study, do not have exclusive drowsiness labels in their hypnograms. Therefore, new datasets and recording setups are required to evaluate this hypothesis.

### 6.2. Kalman Filter versus Kalman Smoother

In this study, the data were processed offline. Therefore, the fixed-interval Kalman smoother was used for tracking the AR model poles. For online applications, a fixed-lag Kalman smoother of an order of 30 s of delay can be used instead, which at the same time benefits from the advantages of smoothing and online analysis.

## 7. Conclusion & Future Work

In this work a robust method was proposed for tracking the poles of a second-order TVAR model for instantaneous frequency estimation of EEG signals. The performance of the Kalman Smoother based approach for accurate IF tracking in low analytical signal envelopes was compared with the conventional Hilbert transform based method. The instantaneous envelope was considered as a complementary feature for the IF and used for automatic sleep stage scoring. The IE and IF features were extracted from simultaneous normalized bandpassed brain rhythms of two-channel sleep EEG signals using the proposed algorithm. The results show that the single feature of IF of beta wave is the most informative among all features for consciousness level detection. The combination of features were used in three different classification methods. The result show the outperformance of the proposed method, in terms of mean accuracy using the LDA classifier. Considering the dynamic nature of sleep stages (temporal evolution from one stage of sleep to another), we expect the classification process to be improved using methods such as the Hidden Markov Model (HMM) and its extensions, which can intrinsically consider the temporal evolution of sleep states.

The utilized leave one subject out cross-validation strategy has made the results subject independent. The compatibility of extracted features with LDA classification led to higher individual performance of subjects in terms of higher overall mean and lower standard deviation of the DA measure (as compared with previous research). Thus, the proposed algorithm could be a reliable approach for automatic sleep staging and a basis for further research.

Studying the *oscillatory instantaneous poles* of high spatial resolution EEG is another future direction of research. As discussed in this work, the concept of drowsiness detection is a possible application of this study, which may lead to a robust alertness detection algorithm with various real-world applications.

## Footnotes

1 In writing equations (13) to (21), simplifications have been made due to the first-order AR dynamics assumed in (10).

## References

[1] K. Šušmáková, “Human sleep and sleep EEG,” Measurement Science Review, vol. 4, no. 2, pp. 59–74, 2004.

[2] (2016) What is Sleep? - American Sleep Association. [Online]. Available: https://www.sleepassociation.org/patients-general-public/what-is-sleep/

[3] A. Rechtschaffen, “A manual of standardized terminology, techniques and scoring system for sleep stages of human subjects,” Public health service, 1968.

[4] D. Moser, P. Anderer, G. Gruber, S. Parapatics, E. Loretz, M. Boeck, G. Kloesch, E. Heller, A. Schmidt, H. Danker-Hopfe et al., “Sleep classification according to AASM and Rechtschaffen & Kales: effects on sleep scoring parameters,” Sleep, vol. 32, no. 2, pp. 139–149, 2009.

[5] L. Doroshenkov, V. Konyshev, and S. Selishchev, “Classification of human sleep stages based on EEG processing using hidden Markov models,” Biomedical Engineering, vol. 41, no. 1, pp. 25–28, 2007.

[6] L. Zoubek, S. Charbonnier, S. Lesecq, A. Buguet, and F. Chapotot, “Feature selection for sleep/wake stages classification using data driven methods,” Biomedical Signal Processing and Control, vol. 2, no. 3, pp. 171–179, 2007.

[7] A. Subasi, “EEG signal classification using wavelet feature extraction and a mixture of expert model,” Expert Systems with Applications, vol. 32, no. 4, pp. 1084–1093, 2007.

[8] F. Ebrahimi, M. Mikaeili, E. Estrada, and H. Nazeran, “Automatic sleep stage classification based on EEG signals by using neural networks and wavelet packet coefficients,” in 2008 30th Annual International Conference of the IEEE Engineering in Medicine and Biology Society. IEEE, 2008, pp. 1151–1154.

[9] R. K. Sinha, “Artificial neural network and wavelet based automated detection of sleep spindles, REM sleep and wake states,” Journal of medical systems, vol. 32, no. 4, pp. 291–299, 2008.

[10] E. Oropesa, H. L. Cycon, and M. Jobert, “Sleep stage classification using wavelet transform and neural network,” International computer science institute, 1999.

[11] L. Fraiwan, K. Lweesy, N. Khasawneh, H. Wenz, and H. Dickhaus, “Automated sleep stage identification system based on time–frequency analysis of a single EEG channel and random forest classifier,” Computer methods and programs in biomedicine, vol. 108, no. 1, pp. 10–19, 2012.

[12] K. Šušmáková and A. Krakovská, “Discrimination ability of individual measures used in sleep stages classification,” Artificial Intelligence in Medicine, vol. 44, no. 3, pp. 261–277, 2008.

[13] A. Flexer, G. Gruber, and G. Dorffner, “A reliable probabilistic sleep stager based on a single EEG signal,” Artificial Intelligence in Medicine, vol. 33, no. 3, pp. 199–207, 2005.

[14] S.-T. Pan, C.-E. Kuo, J.-H. Zeng, and S.-F. Liang, “A transition-constrained discrete hidden Markov model for automatic sleep staging,” Biomedical engineering online, vol. 11, no. 1, p. 1, 2012.

[15] S. Özşen, “Classification of sleep stages using class-dependent sequential feature selection and artificial neural network,” Neural Computing and Applications, vol. 23, no. 5, pp. 1239–1250, 2013.

[16] F. Chapotot and G. Becq, “Automated sleep–wake staging combining robust feature extraction, artificial neural network classification, and flexible decision rules,” International Journal of Adaptive Control and Signal Processing, vol. 24, no. 5, pp. 409–423, 2010.

[17] A. Subasi, M. K. Kiymik, M. Akin, and O. Erogul, “Automatic recognition of vigilance state by using a wavelet-based artificial neural network,” Neural Computing & Applications, vol. 14, no. 1, pp. 45–55, 2005.

[18] M. E. Tagluk, N. Sezgin, and M. Akin, “Estimation of sleep stages by an artificial neural network employing EEG, EMG and EOG,” Journal of medical systems, vol. 34, no. 4, pp. 717–725, 2010.

[19] M. Ronzhina, O. Janoušek, J. Kolářová, M. Nováková, P. Honzík, and I. Provazník, “Sleep scoring using artificial neural networks,” Sleep medicine reviews, vol. 16, no. 3, pp. 251–263, 2012.

[20] B. Şen, M. Peker, A. Çavuşoğlu, and F. V. Çelebi, “A comparative study on classification of sleep stage based on EEG signals using feature selection and classification algorithms,” Journal of medical systems, vol. 38, no. 3, pp. 1–21, 2014.

[21] S. Gudmundsson, T. P. Runarsson, and S. Sigurdsson, “Automatic sleep staging using support vector machines with posterior probability estimates,” in International Conference on Computational Intelligence for Modelling, Control and Automation and International Conference on Intelligent Agents, Web Technologies and Internet Commerce (CIMCA-IAWTIC’06), vol. 2. IEEE, 2005, pp. 366–372.

[22] B. Koley and D. Dey, “An ensemble system for automatic sleep stage classification using single channel EEG signal,” Computers in biology and medicine, vol. 42, no. 12, pp. 1186–1195, 2012.

[23] M. V. Yeo, X. Li, K. Shen, and E. P. Wilder-Smith, “Can SVM be used for automatic EEG detection of drowsiness during car driving?” Safety Science, vol. 47, no. 1, pp. 115–124, 2009.

[24] Y.-H. Lee, Y.-S. Chen, and L.-F. Chen, “Automated sleep staging using single EEG channel for REM sleep deprivation,” in Bioinformatics and BioEngineering, 2009. BIBE’09. Ninth IEEE International Conference on. IEEE, 2009, pp. 439–442.

[25] T. Lajnef, S. Chaibi, P. Ruby, P.-E. Aguera, J.-B. Eichenlaub, M. Samet, A. Kachouri, and K. Jerbi, “Learning machines and sleeping brains: automatic sleep stage classification using decision-tree multi-class support vector machines,” Journal of neuroscience methods, vol. 250, pp. 94–105, 2015.

[26] S. Motamedi-Fakhr, M. Moshrefi-Torbati, M. Hill, C. M. Hill, and P. R. White, “Signal processing techniques applied to human sleep EEG signals—A review,” Biomedical Signal Processing and Control, vol. 10, pp. 21–33, 2014.

[27] L. Cohen, Time-frequency analysis. Prentice hall, 1995, vol. 778.

[28] L. Patomäki, J. Kaipio, and P. A. Karjalainen, “Tracking of nonstationary EEG with the roots of ARMA models,” in IEEE Conf. EMBC, vol. 95, 1995.

[29] Z. Rogowski, I. Gath, and E. Bental, “On the prediction of epileptic seizures,” Biological cybernetics, vol. 42, no. 1, pp. 9–15, 1981.

[30] N. Dahal, D. N. Nandagopal, B. Cocks, R. Vijayalakshmi, N. Dasari, and P. Gaertner, “TVAR modeling of EEG to detect audio distraction during simulated driving,” Journal of neural engineering, vol. 11, no. 3, p. 036012, 2014.

[31] A. Lashkari, R. Boostani, and S. Afrasiabi, “Estimation of the anesthetic depth based on instantaneous frequency of electroencephalogram,” in Telecommunications and Signal Processing (TSP), 2015 38th International Conference on. IEEE, 2015, pp. 403–407.

[32] D. P. Nguyen, M. A. Wilson, E. N. Brown, and R. Barbieri, “Measuring instantaneous frequency of local field potential oscillations using the Kalman smoother,” Journal of neuroscience methods, vol. 184, no. 2, pp. 365–374, 2009.

[33] B. Picinbono, “On instantaneous amplitude and phase of signals,” IEEE Transactions on signal processing, vol. 45, no. 3, pp. 552–560, 1997.

[34] R. Sameni and E. Seraj, “A robust statistical framework for instantaneous electroencephalogram phase and frequency estimation and analysis,” Physiological measurement, vol. 38, no. 12, p. 2141, 2017.

[35] B. Boashash, “Estimating and interpreting the instantaneous frequency of a signal. II. Algorithms and applications,” Proceedings of the IEEE, vol. 80, no. 4, pp. 540–568, 1992.

[36] L. B. Almeida, “The fractional Fourier transform and time-frequency representations,” IEEE Transactions on signal processing, vol. 42, no. 11, pp. 3084–3091, 1994.

[37] H. K. Kwok and D. L. Jones, “Improved instantaneous frequency estimation using an adaptive short-time Fourier transform,” IEEE Transactions on Signal Processing, vol. 48, no. 10, pp. 2964–2972, 2000.

[38] B. Boashash, “Estimating and interpreting the instantaneous frequency of a signal. I. Fundamentals,” Proceedings of the IEEE, vol. 80, no. 4, pp. 520–538, 1992.

[39] Y. Grenier, “Time-dependent ARMA modeling of nonstationary signals,” IEEE Transactions on Acoustics, Speech, and Signal Processing, vol. 31, no. 4, pp. 899–911, 1983.

[40] G. E. Box, G. M. Jenkins, G. C. Reinsel, and G. M. Ljung, Time series analysis: forecasting and control. John Wiley & Sons, 2015.

[41] M. Arnold, X. Milner, H. Witte, R. Bauer, and C. Braun, “Adaptive AR modeling of nonstationary time series by means of Kalman filtering,” IEEE Transactions on Biomedical Engineering, vol. 45, no. 5, pp. 553–562, 1998.

[42] M. Aboy, O. W. Márquez, J. McNames, R. Hornero, T. Trong, and B. Goldstein, “Adaptive modeling and spectral estimation of nonstationary biomedical signals based on Kalman filtering,” IEEE transactions on biomedical engineering, vol. 52, no. 8, pp. 1485–1489, 2005.

[43] S. M. Kay, Fundamentals of statistical signal processing, volume I: estimation theory. Prentice Hall, 1993.

[44] E. R. Dougherty, Random processes for image and signal processing. SPIE Optical Engineering Press, 1999.

[45] M. P. Tarvainen, J. K. Hiltunen, P. O. Ranta-aho, and P. A. Karjalainen, “Estimation of nonstationary EEG with Kalman smoother approach: an application to event-related synchronization (ERS),” IEEE Transactions on Biomedical Engineering, vol. 51, no. 3, pp. 516–524, 2004.

[46] S. Haykin, “Adaptive filter theory,” Englewood Cliffs, NJ, Prentice-Hall, 1986. 607, 1986.

[47] R. G. Brown, Introduction to random signal analysis and Kalman filtering. John Wiley & Sons, 1983.

[48] A. Gelb, Applied optimal estimation. MIT press, 1974, sixteenth print 2001.

[49] R. Sameni, M. B. Shamsollahi, C. Jutten, and G. D. Clifford, “A nonlinear Bayesian filtering framework for ECG denoising,” IEEE Transactions on Biomedical Engineering, vol. 54, no. 12, pp. 2172–2185, 2007.

[50] M. A. Maasoumnia, “Estimation Theory and Optimal Filtering, [Lecture Notes],” 2003, Sharif University of Technology, Tehran, Iran.

[51] R. Sameni, “A linear kalman notch filter for power-line interference cancellation,” in Artificial Intelligence and Signal Processing (AISP), 2012 16th CSI International Symposium on. IEEE, 2012, pp. 604–610.

[52] B. Kemp, A. H. Zwinderman, B. Tuk, H. A. Kamphuisen, and J. J. Oberye, “Analysis of a sleep-dependent neuronal feedback loop: the slow-wave microcontinuity of the EEG,” IEEE Transactions on Biomedical Engineering, vol. 47, no. 9, pp. 1185–1194, 2000.

[53] B. Van Sweden, B. Kemp, H. Kamphuisen, and A. Ver der Velde, “Alternative electrode placement in (automatic) sleep scoring,” Sleep, vol. 13, no. 3, pp. 279–283, 1990.

[54] J. C. Armington and L. L. Mitnick, “Electroencephalogram and sleep deprivation,” Journal of applied physiology, vol. 14, no. 2, pp. 247–250, 1959.

